# Sensitivity analysis of models of gas exchange for lung hyperpolarised 129-Xe MRS and MRI

**DOI:** 10.1101/2023.07.26.550733

**Authors:** Yohn Taylor, Frederick J Wilson, Mina Kim, Geoff J M Parker

**Author notes:** Corresponding author: **Name** Geoff J M Parker, **Department** Centre for Medical Image Computing, **Institute** University College London, **Address** 90 High Holborn, WC1V 6LJ, London, United Kingdom, **E-mail**.

## Abstract

**Purpose:** Sensitivity analysis enables the identification of influential parameters and the optimisation of model composition. Such methods have not previously been applied systematically to models describing hyperpolarised ^129^Xe gas exchange in the lung. Here, we evaluate current ^129^Xe gas exchange models to assess their precision for identifying alterations in pulmonary-vascular function and lung microstructure.

**Methods:** We assess sensitivity using established univariate methods and scatter plots for parameter interactions. We apply them to the model described by Patz and MOXE *et al*., examining their ability to measure: i) importance (rank), ii) temporal dependence, and iii) interaction effects of each parameter across healthy and diseased ranges.

**Results:** The univariate methods and scatter plot analyses demonstrate consistently similar results for the importance of parameters common to both models evaluated. Alveolar surface area to volume ratio is identified as the parameter to which model signals are most sensitive. The alveolar-capillary barrier thickness is identified as a low-sensitivity parameter for the MOXE model. An acquisition window of at least 200 ms effectively demonstrates model sensitivity to most parameters. Scatter plots reveal interaction effects in both models, impacting output variability and sensitivity.

**Conclusion:** Our sensitivity analysis ranks the parameters within the model described by Patz *et al* and within the MOXE model. The MOXE model shows low sensitivity to alveolar-capillary barrier thickness, highlighting the need for designing acquisition protocols optimised for the measurement of this parameter. The presence of parameter interaction effects highlights the requirement for care in interpreting model outputs.

## Introduction

Hyperpolarised ^129^Xe (HpXe) MRS and MRI of the lungs permit the assessment of the temporal dynamics of gas exchange across the alveolar-capillary barrier (1). ^129^Xe is highly lipophilic and soluble and follows a similar diffusion pathway to that of oxygen (9). As a result, ^129^Xe gas transfer mechanisms can be modelled utilising uptake curves of characteristic spectroscopic peaks generated from the movement of ^129^Xe between the gas (airways and alveoli) and dissolved (tissue/blood) lung compartments. Fitting 1D diffusion compartment models to HpXe MRS chemical shift saturation recovery (CSSR) data enables the extraction of microstructural and physiological measurements of the lung, providing markers for evaluating global and regional lung function. Two-compartment models (11; 16) typically consider ^129^Xe dissolved in blood and tissue collectively - the dissolved phase - while three-compartment models (8; 4) differentiate between frequency shifted ^129^Xe peaks from a combined tissue and plasma pool - the tissue phase - and red blood cells (RBC). The two-compartment model introduced by Patz *et al* (11) (from here on denoted ‘Patz model’), and the three-compartment MOXE model (4) have been used for the clinical assessment of a range of lung diseases, including chronic obstructive pulmonary disease (COPD), interstitial lung disease (ILD) and asthma (14; 7).

The Patz model allows the estimation of septal thickness, *d*, the alveolar surface area to volume ratio, SVR, and the blood transit time through the capillary bed, τ_*c*_. As an extension of the Patz model, the MOXE model additionally considers the tissue and RBC phases (11), allowing the estimation of three more parameters: the alveolar-capillary barrier thickness, *δ*, the xenon-exchange time constant, *T*, and the haematocrit, HCT.

When using either model to characterise disease it is necessary to understand the attainable measurement specificity and sensitivity to underlying disease processes for each parameter. The type and severity of disease will dictate the parameters that are expected to provide disease sensitivity. For example, a decrease in SVR and an increase in *d* are indicative of symptoms common within emphysematic and fibrotic lung diseases (5), respectively. The ability of a measurement technique and an associated signal model to precisely measure these differences is dependent on the dynamic range of signal changes that are associated with changes in the model’s parameters. Sensitivity analysis investigates how variations in signals predicted by a model can be attributed to variations within different input factors/parameters (18). If model output is not sensitive to changes within the relevant parameter space for a particular experimental setting, precise interrogation of these parameters cannot be achieved. As a result, sensitivity analysis can assist in defining influential and non-influential parameters within a given model, thereby identifying what parameterisations are likely to be feasible and informative within a given experimental scenario (20).

In this study we apply both univariate and multivariate sensitivity analysis methods to the Patz and MOXE 1D diffusion models to investigate each model’s sensitivity to parameter variability. We identify interaction effects, determine parameter importance - ranking parameters to which the output is most and least sensitive to (12) - for both models, and identify model sensitivity to parameter variation across a range of saturation recovery repetition times; identifying experimental periods for optimal parameter measurement.

## Theory

### Chemical shift saturation recovery (CSSR)

The rate of regrowth due to the influx of polarised spins of the ^129^Xe dissolved phase signal after a saturation radio-frequency (RF) pulse can be described by appropriate models, allowing parameters of interest to be estimated. When using chemical shift saturation recovery (CSSR) (2; 11), a spectroscopic measurement of the ^129^Xe magnetisation is taken after each of a series of repetition times TR, allowing the rate of ^129^Xe uptake within the dissolved phase compartments to be determined. Uptake curves demonstrate a characteristic increase over time, highlighting an initial rapid influx of gas within the compartments, followed by a slower uptake phase. This can be represented by a combination of two processes: the replenishment of the ^129^Xe from the gas phase to the tissue phase, and the uptake into and exit from the local tissue of ^129^Xe via the capillary blood flow.

### Two compartment model

The model described by Patz (11) is derived from the solution of the diffusion equation,

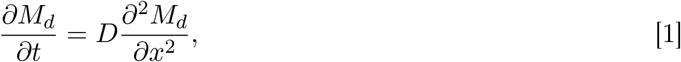

where *D* denotes the dissolved phase diffusion coefficient of xenon. The diffusion process is described as a two compartment model, with the septum bounded by the alveoli as shown in Fig. 1. The Patz model describes the diffusion of hyperpolarised ^129^Xe into the dissolved phase, defined as a single compartment combining the tissue phase and red blood cells.

**Figure 1.**
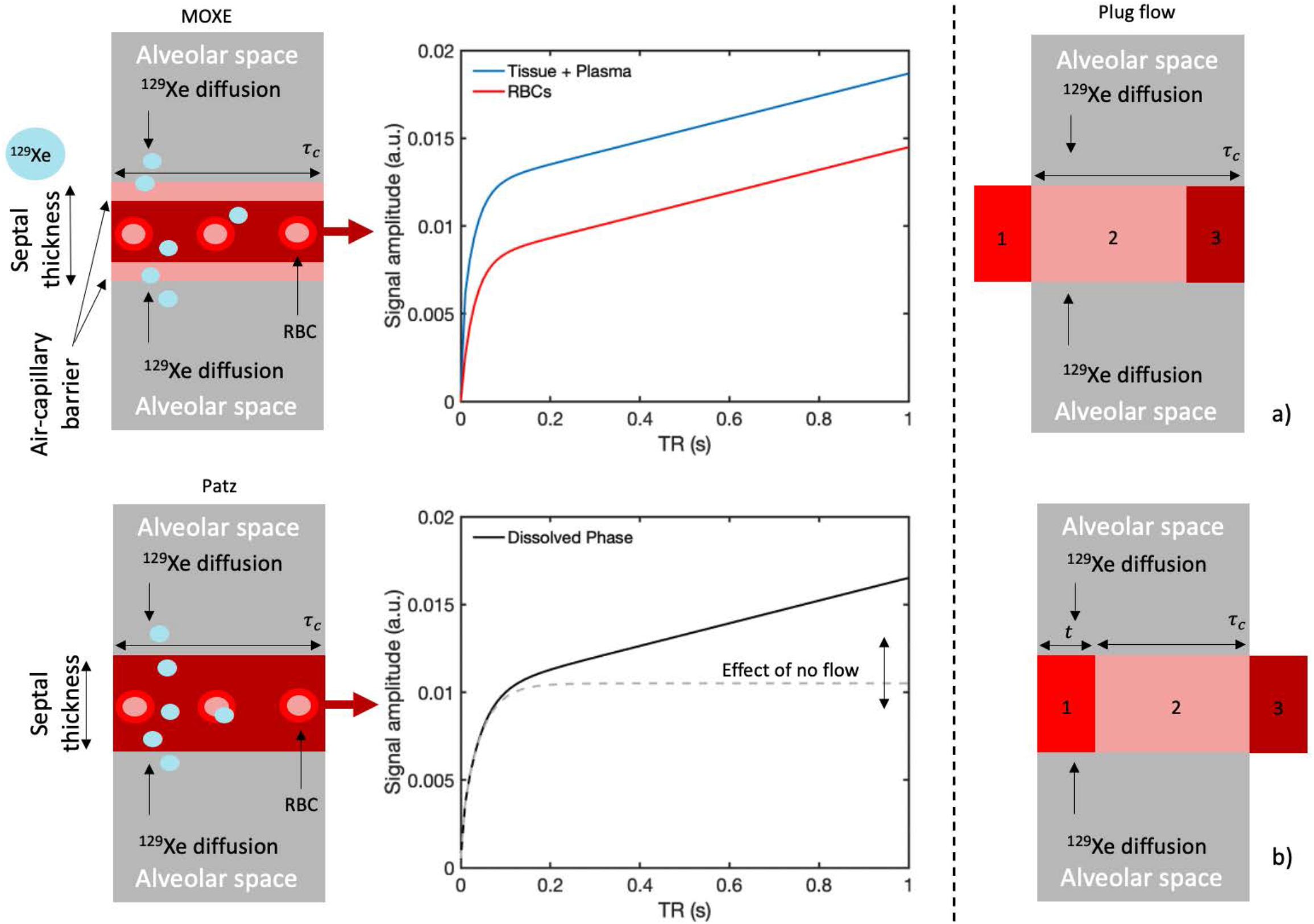
Block diagrams of the MOXE and Patz diffusion systems with ^129^Xe diffusing from both boundaries alongside corresponding model-derived signal plots. The Patz model is represented by a single plot of the dissolved phase signal, whereas the MOXE model is a coupled equation containing plots for both the tissue and RBC compartments. Plug flow diagrams (a and b) highlight defined spatial regions of blood quantifying blood flow.

When employing the CSSR technique, hyperpolarised ^129^Xe is inhaled into the lungs followed by a frequency selective RF pulse, which saturates the longitudinal magnetisation in the dissolved phase. At time t = 0, *M*_*g*_(x, t = 0) = *M*_0_ and *M*_*d*_(*x, t* = 0) = 0, where *x* represents the spatial position of dissolved xenon and *M*_*g*_ and *M*_*d*_ represent the longitudinal magnetisation in the gas phase (alveoli) and dissolved phase (septum), respectively. At *t >* 0, unsaturated ^129^Xe from the alveoli diffuses into the septum, leading to

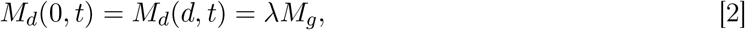

where λ is the Ostwald solubility of xenon within tissue and the magnetisation from the alveoli space *M*_*g*_ is approximated to be unchanged *M*_*g*_(*x, t*)= *M*_0_. The solution for the boundary conditions for ^129^Xe magnetisation, *M*_*d*_, within the dissolved phase (2) results in,

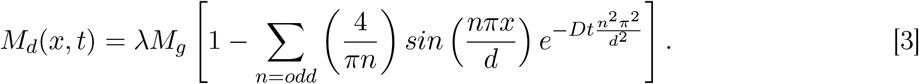

The integral of the ^129^Xe magnetisation in the septum (dissolved phase) provides the fraction of the septum occupied by hyperpolarised ^129^Xe at time *t*,

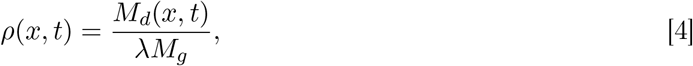

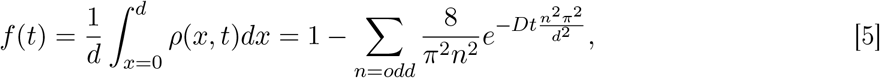

where *ρ* is the dimensionless representation of ^129^Xe magnetisation density within the septal slab of thickness *d*. The ratio of the dissolved phase ^129^Xe signal after time *t* to the gas phase at time *t* =0 is

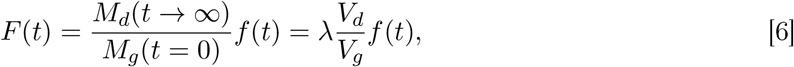

where *M*_*d*_(*t* ⟶ ∞) is the ^129^Xe magnetisation at septum saturation (19). The septal volume is denoted by *V*_*d*_ = *Ad*, where *A* is the surface area between the alveolar gas and the septal boundary. As there are two boundaries, the total surface area *S*_*A*_ = 2*A*. Consequently, the septal volume *V*_*d*_ is

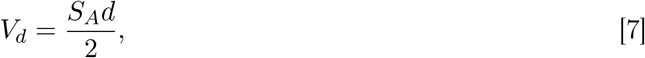

and

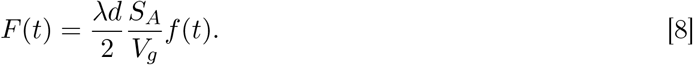

Blood flow is assumed to have constant velocity, to be independent of ^129^Xe concentration, and to be orthogonal to the capillary wall. The Patz model treats the entire septum as a flowing system, with blood flowing within the pulmonary vessels residing in the gas exchange zone (GEZ) for a certain period of time. A simple plug flow model is used in simulating this effect by separating the blood into three spatial regions as shown in Fig. 1.

In Fig. 1, region 1 represents blood upstream from the GEZ at *t* = 0, which subsequently passes through the GEZ for a fraction of the measurement time. Region 2 depicts blood present within the GEZ for the entire duration of *t*. Similarly, region 3 like region 1 spends a fraction of time within the GEZ and is found downstream of the GEZ at time *t*. The figure shows the transit time of the dissolved phase within the GEZ, with panels 1a) and 1b) illustrating the plug flow regions at *t* =0 and *t*, respectively. Due to region 2 residing in the GEZ for the entire diffusion time, the fraction of blood within the septum containing ^129^Xe is *f*(*t*). The fraction of the alveolar space contributing to the diffusion of ^129^Xe into the blood within region 2 is *f*_2_ = (τ_*c*_ *-t*)*/* τ_*c*_.

The fraction of septal space contributing to blood in region 1 and region 3 (*f*_1_, *f*_3_) occupied by magnetised ^129^Xe, differs from region 2 due to their respective starting and ending positions residing outside of the GEZ. Consequently, the incremental time *t*^*′*^ of the diffusion time *t* of region 1 and 3 within the GEZ is used to determine the average *f*_1_, *f*_3_:

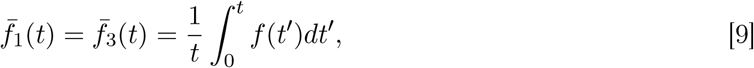

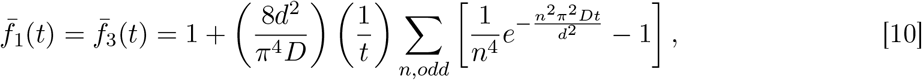

with *f*_3_ = *f*_1_ = *t/*τ_*c*_. The contribution of each region to the overall blood flow is denoted by *F*_1_(*t*) and *F*_3_(*t*):

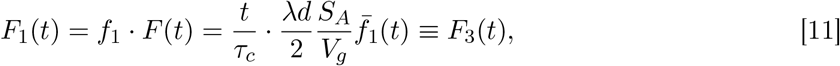

with the total flow given by

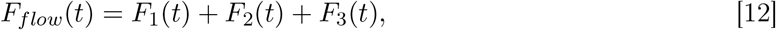

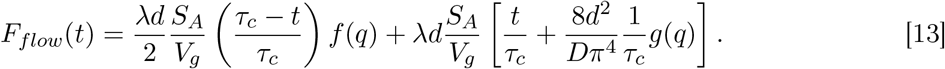

The functions of the dimensionless parameter *q, h*(*q*) and *g(q)* (where *q* = (Dt/d^2^)) are:

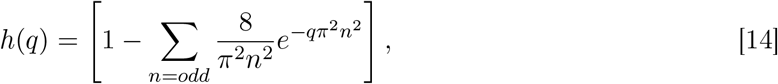

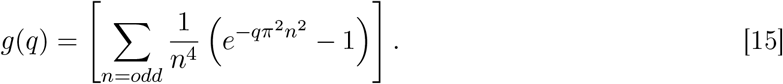

The physiological parameters extracted when utilising the Patz model are the septal thickness *d*, the alveolar surface area to volume of gas ratio SVR (*S*_*A*_*/V*_*g*_) and the capillary transit time, τ_*c*_.

### Three compartment model

#### MOXE model

The MOXE model is an extension of the Patz model, incorporating both tissue and RBC dynamics (4). As a result, the alveolar-capillary barrier thickness, *δ*, exchange time constant, *T*, and the haematocrit, HCT, can be derived, with the introduction of terms for the relative Ostwald solubilities of plasma and RBC (*λ*_*PL*_ and *λ*_*RBC*_). The signal distribution *S*_*d*_ is equivalent to the ratio of dissolved and gas phase magnetisation, *F* (Eq. 8) such that

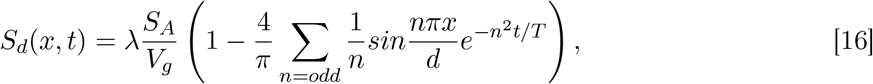

where *T*, is the exchange time constant represented by

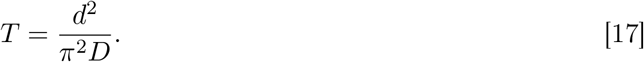

The spatial integral of *S*_*d*_ over 0 to *δ* and *d - δ* to *d*, as highlighted in Fig. 1, leads to the signal contribution *S*_*d*1_ of the ^129^Xe signal within the tissue phase:

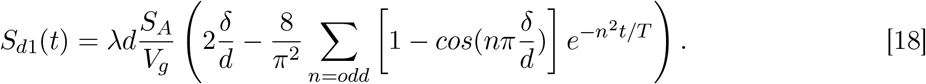

Modelling the ^129^Xe signal from the blood, incorporating flow, results in

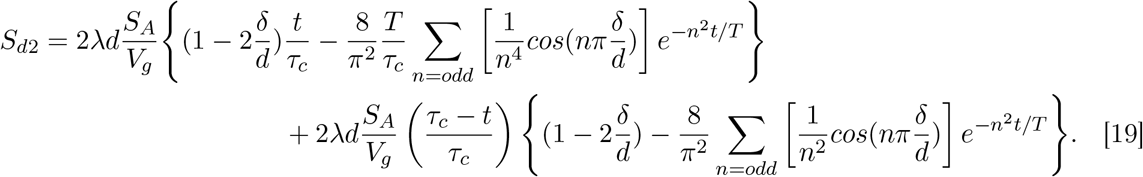

Expressions for the signal amplitudes of the chemical shifts within the dissolved tissue and RBCs displayed in Fig. 1 are given by

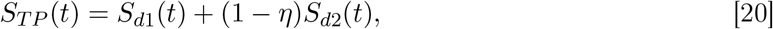

and

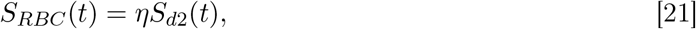

where *η* is the fraction of dissolved gas in the RBC’s and (1 *-η*)*S*_*d*2_(*t*) is the term for the signal contribution within the plasma, with HCT defined as

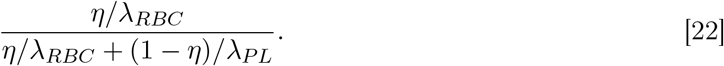

The plots for the Patz and MOXE models in Fig. 1 both display a rapid signal increase until *t* ∼ 100 ms due to the influx of ^129^Xe within the septum. After this point, contribution to signal growth is exclusively a result of blood flow and increases more slowly with time. Fig. 1b highlights the additional signal change in the Patz model when capillary transit time τ_*c*_ increases to values greater than the diffusion time due to blood flow. As τ_*c*_ → ∞ (i.e. as blood flow tends to zero), the ^129^Xe signal increase after *t* ∼ 100 ms becomes negligible; as a result, ^129^Xe saturation occurs, highlighted in Fig. 1b by the dashed line. A similar phenomenon is seen with the MOXE model, but is not shown here to retain diagram clarity.

## Methods

Univariate and multivariate analysis (13) methods were applied in the assessment of each parameter for each model, with values spanning ranges that include expected values for both healthy and diseased lungs, as shown in Table. 1.

### Univariate sensitivity analysis

Model parameter range was defined as the mean *±* standard deviation of the literature-derived parameter values (Table. 1). Parameters were independently varied within these ranges, whilst maintaining constant values for the other parameters at their mean.

Python 3 (spyder 5.0.0) and MATLAB (2021a) were used for the CSSR Patz and MOXE model simulations and sensitivity analysis implementation. CSSR simulations with forty TR values equally spaced between 10 ms and 800 ms were generated for each model’s univariate sensitivity analysis plot. Numerical computations of the infinite series of *F*_*flow*_, *S*_*d*1_, and *S*_*d*2_ were reduced at the fifth terms, removing values of *n* =9 or greater.

The sensitivity of each model to its constituent parameters was simulated by incrementing each parameter independently over its range at each TR whilst maintaining the other parameters at their mean value. This was then visualised by plotting the resultant signal range as a function of TR.

### Signal Percent Change

Signal percentage change (SPC) refers to the percentage signal change observed as a parameter is varied from its mean value, simulated for each of the forty TRs. Simulations employed the upper and lower limits of a specific parameter’s range whilst maintaining the other parameters at their mean values. Simulations outline the general shape of signal change due to variation over the parameter range and at which point in the experimental procedure this change occurs; determining if specific periods within the acquisition window exhibit greater changes in signal than others. Constant SPC displays a lack of variation in the sensitivity within the experimental procedure whilst any change in signal is associated with a change in sensitivity.

### Scatter plots sensitivity analysis

Scatter plots are commonly used as visual methods for examining correlations between model output signal variability and input parameters. In general, parameters producing a larger degree of output variability - displaying visible negative and positive correlations - are parameters to which the model is more sensitive. 2D plots were produced by simulating all the output values (across all values of other parameters) corresponding to each individual parameter value. 3D plots demonstrate the relationship between each parameter and signal output when also considering the range of a second chosen parameter, allowing evidence for parameter interaction to be observed.

Latin hypercube sampling was used to efficiently sample the parameter space evenly for parameters in both models. All scatter simulations and sampling were performed using both custom-written and built-in MATLAB code (The Mathworks, Natick MA, R2021a).

## Results

### Univariate sensitivity analysis

The univariate sensitivity analysis for each parameter of the Patz model is shown in Fig. 2. The three plots highlight the signal variations observed when independently varying each model parameter SVR, *d*, τ_*c*_. The colour scale corresponds to the values of the parameter being varied. For all parameters, an increase in model output signal amplitude and signal range is observed when moving to longer TRs. As shown by the gradations in the colour coding, increases in both SVR and *d* lead to increased output signal amplitude, whereas increases in τ_*c*_ lead to a decrease in output signal amplitude. A larger model output signal range for SVR than for *d* or τ_*c*_ is apparent at all TRs, suggesting higher signal sensitivity to this model parameter.

**Figure 2.**
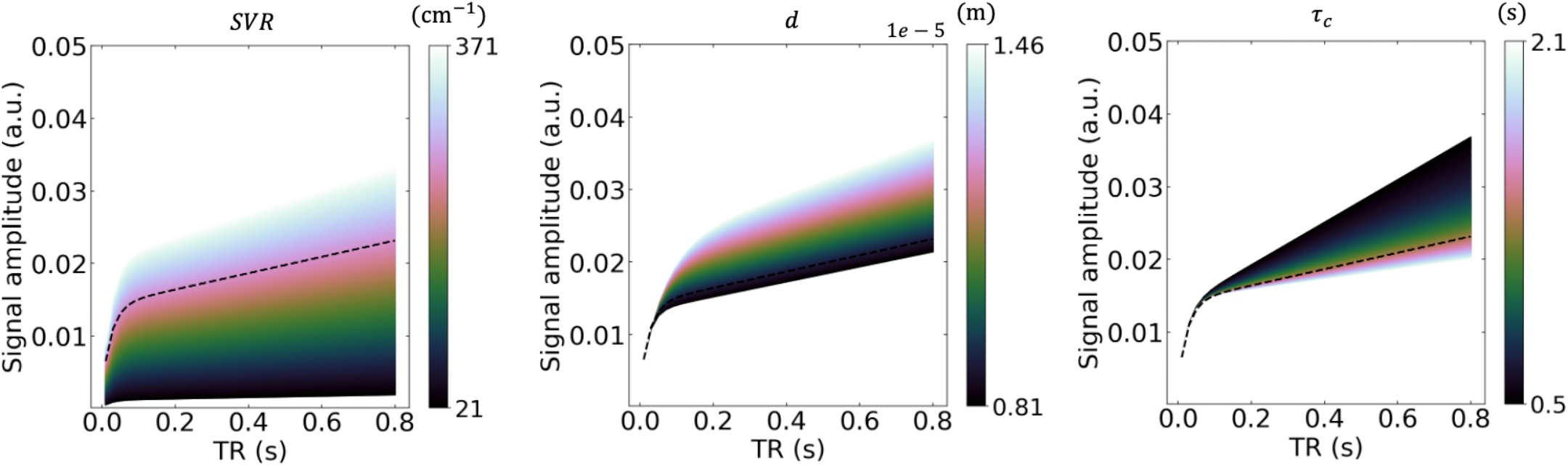
Univariate sensitivity analysis plots of the Patz parameters spanning the allowed range, where the black dashed lines are the outputs associated with input Patz model values for a healthy individual (Table 1) (11). Colour bars correspond to the defined range of each parameter.

Fig. 3 and Fig. 4 show the univariate sensitivity analysis for the tissue phase and RBC phase of the MOXE model, respectively. Similar results to the Patz model for SVR, *d*, and τ_*c*_ are present in both the tissue phase and RBC plots although, in general, sensitivity to τ_*c*_ is lower for MOXE. Of the three additional MOXE parameters, HCT displays the largest output range of signal intensity for most of the TR range, followed by *T* and then *δ*, when considering the tissue phase. For the RBC phase, a larger output range is generally seen for *δ* than for *T*, with both ranges again being smaller than that for HCT. As with τ_*c*_, increases in HCT lead to decreases in signal amplitude when considering the tissue phase, but this trend is reversed for HCT in the RBC plots. Increases in *T* lead to decreases in signal for both the tissue phase and RBC phase. Increases in *δ* lead to increases in signal amplitude when considering the tissue phase, but to signal decreases in the RBC plots. The range of signal intensity variation due to varying HCT increases with TR, whereas the opposite trend is seen for *δ* and *T* in the tissue phase. In the RBC phase the range of signal intensity due to variation in *δ* increases with TR, with *T* again showing a decrease.

**Table 1:** Symbol, meaning, range, typical healthy values, and units of parameters used within the Patz and MOXE models.

| Symbol | Meaning | Range (mean ± st. dev.) | Healthy | Units (citations) |
| --- | --- | --- | --- | --- |
| $SVR$ | Surface area to volume ratio | $196 \pm 175$ | 256 | $\text{cm}^{-1}$ (5) |
| $d$ | Septal wall thickness | $11.35 \pm 3.25$ | 8.8 | $\mu\text{m}$ (4; 17) |
| $\tau_c$ | Capillary transit time | $1.3 \pm 0.8$ | 1.3 | s (21) |
| $T$ | Exchange time constant | $0.039 \pm 0.026$ | 0.026 | ms (4; 17) |
| $HCT$ | Haematocrit | $0.225 \pm 0.085$ | 0.265 | % (4; 19) |
| $\delta$ | Alveolar-capillary barrier thickness | $1.2065 \pm 0.4445$ | 1.0 | $\mu\text{m}$ (3; 19) |

**Figure 3.**
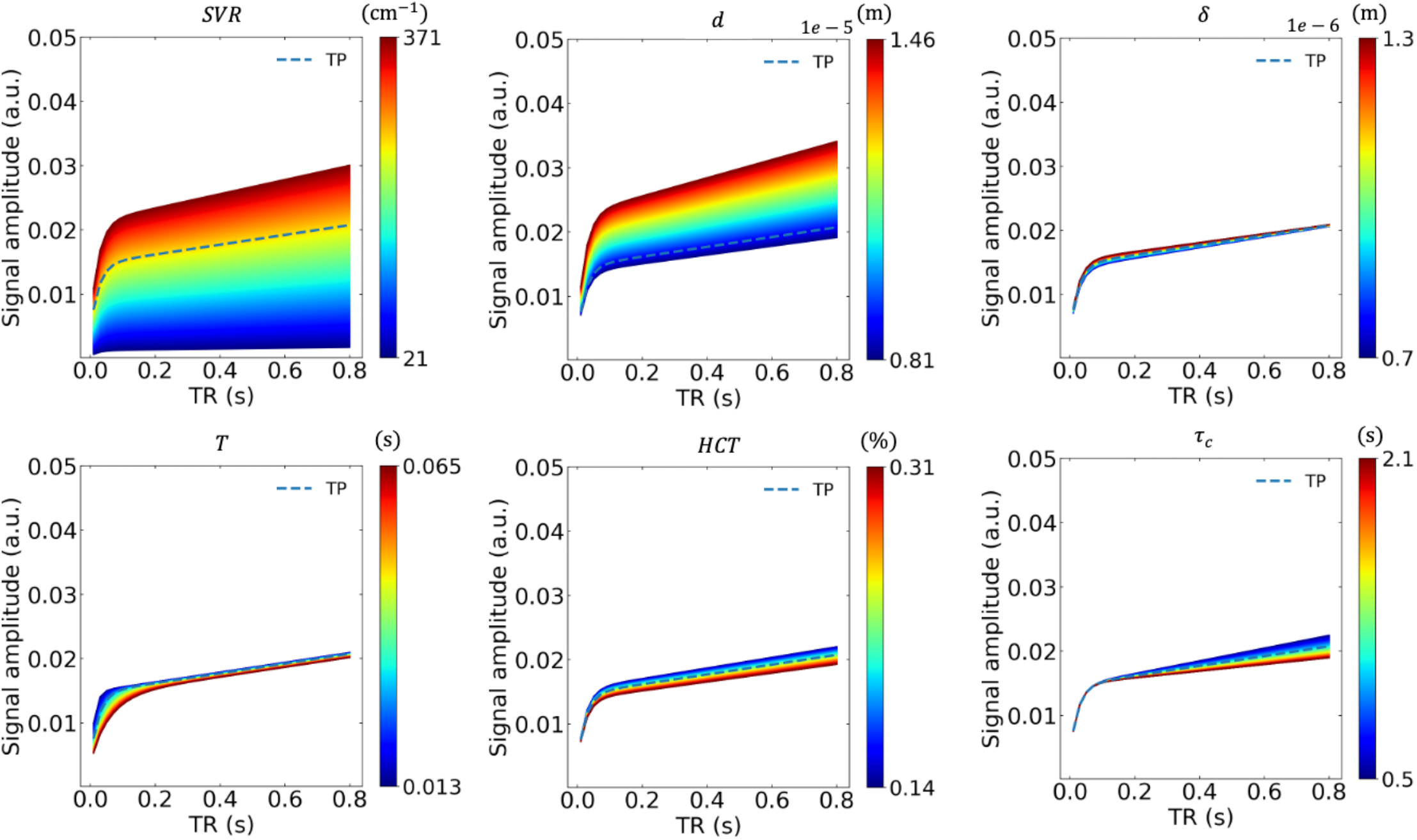
Univariate sensitivity analysis plots of the MOXE parameters spanning the allowed range (tissue phase), where the black dashed lines are the outputs associated with input MOXE model values for a healthy individual (Table 1).

**Figure 4.**
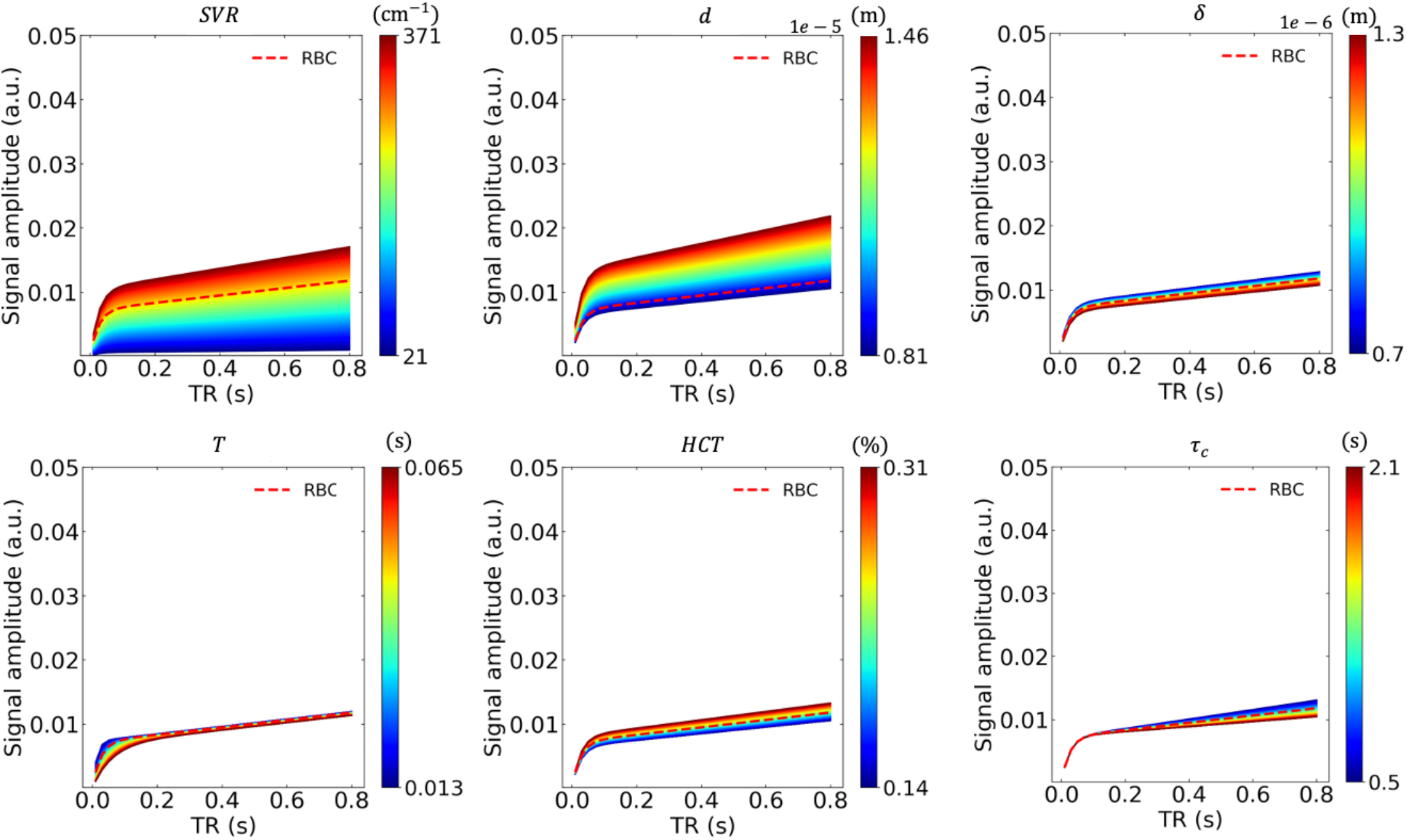
Univariate sensitivity analysis plots of the MOXE parameters spanning the allowed range (RBC phase), where the black dashed lines are the outputs associated with input MOXE model values for a healthy individual (Table 1).

### Signal percentage change analysis

The SPC plots in Fig. 5 display the upper and lower SPC values corresponding to the maximum and minimum parameter values for each Patz and MOXE parameter range throughout the simulated range of TRs. Variations in SPC over the TR range for the tissue phase plots were evident in all parameters with the exception of SVR, whilst variations in SPC for the RBC phase are only evident in d, *δ, T* and τ_*c*_. The largest shift in the shape of SPC is seen between 0 ms *<* TR *<* 200 ms, with the exception of changes due to τ_*c*_, which displays a gradual change throughout the TR range.

**Figure 5.**
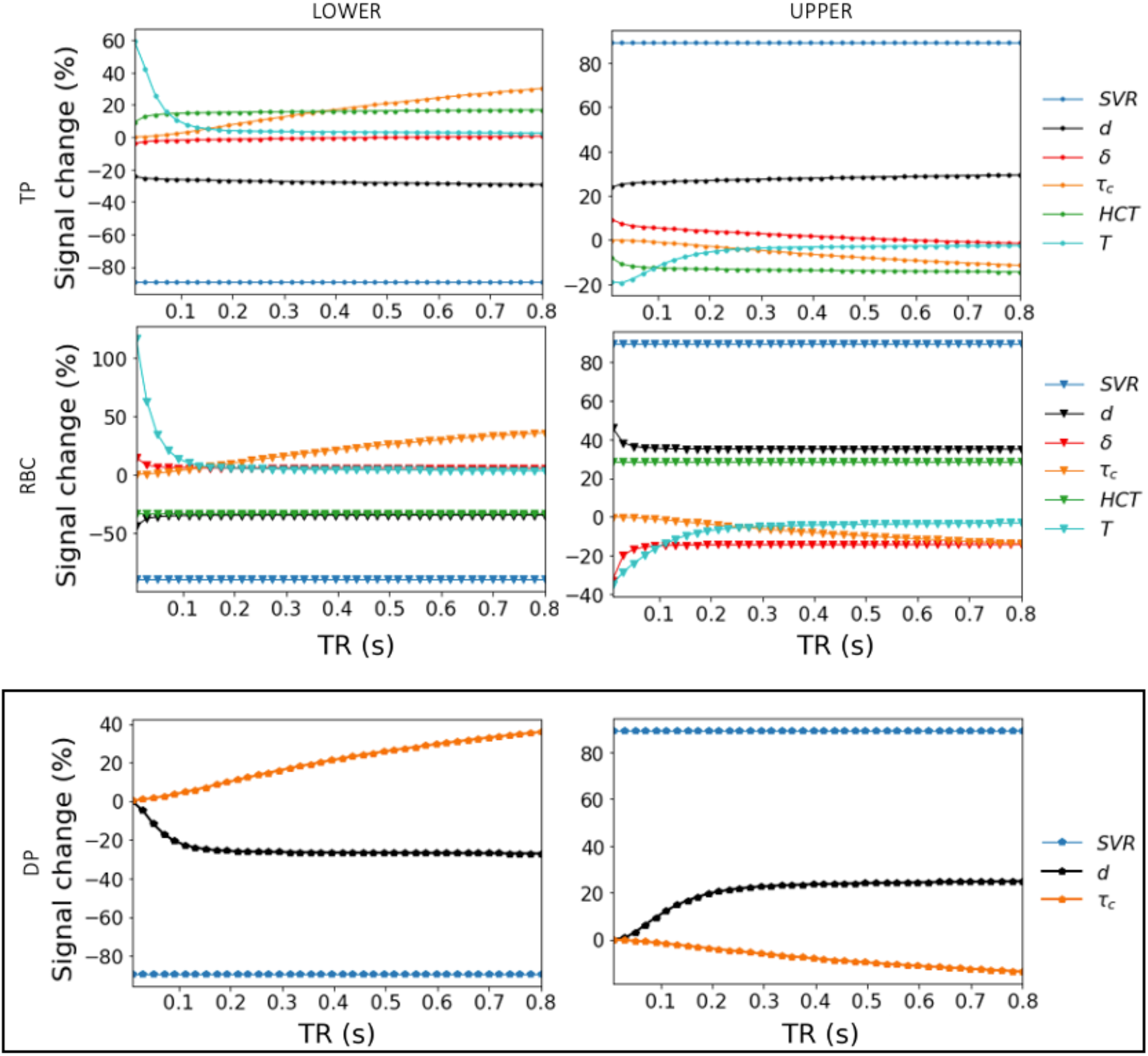
Signal percentage change (SPC) as a function of delay time (TR), highlighting the extremes of the parameter range for the tissue phase (TP) and RBC compartments of the MOXE model (top two rows) and for the Patz model dissolved phase (DP) (bottom row, boxed). The upper and lower limits are the signal percentage change values when using the highest and lowest values within each individual parameter range.

### Scatter plot and 3D plot analyses

Fig. 6 displays scatter plots for the Patz model, showing the signal amplitude change due to each individual parameter when accounting for the full variation of the other model parameters. Plots are shown for simulations using a TR midway between the initial and final TR at ∼ 400 ms. The strongest correlation with signal amplitude is seen for SVR, with τ_*c*_ showing the weakest evidence for correlation, suggesting that distinguishing signal changes uniquely due to τ_*c*_ would be the most challenging. 3D plots in Fig. 7 display parameter interaction effects between the three Patz model parameters with the boxed (red and black) plots (τ_*c*700_) highlighting the interaction effects of using a longer TR.

**Figure 6.**
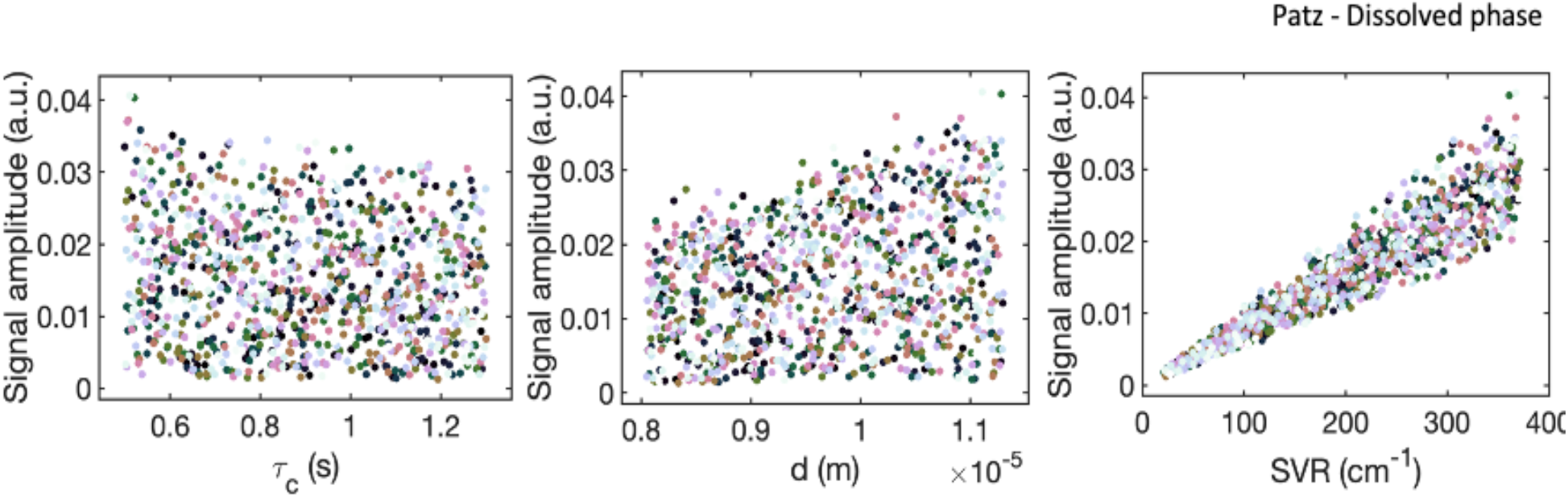
2D Scatter plots showing the signal relationship with each individual parameter whilst allowing the full variation of the other model parameters.

**Figure 7.**
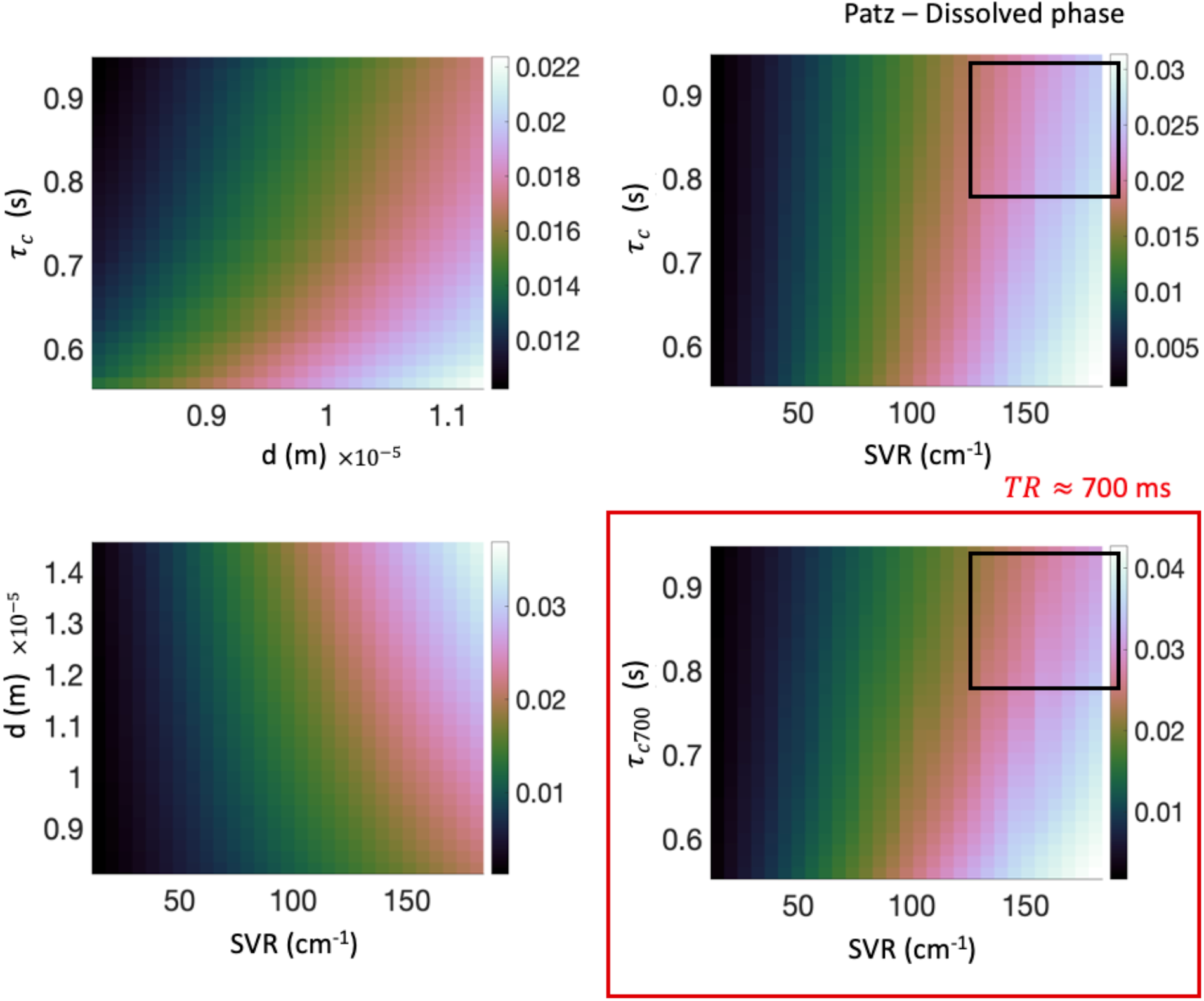
3D plots show the relationship between each parameter considering the range of the other parameters against the signal output. All simulations were taken at a TR of 400 ms with the exception of τ_*c*700_, which was taken at a TR of 700 ms (red box); demonstrating a dependency between interaction effects and TR (black boxes).

Figs. 8 and 9 show the signal amplitude change of each individual parameter when accounting for the full variation of the other model parameters for the MOXE model, for the tissue phase and RBC phase, respectively. Plots are shown for simulations using a TR midway between the initial and final TR at ∼ 400 ms. The strongest correlations at TR = 400 ms are again evident in SVR, with *d* and HCT also showing correlations.

**Figure 8.**
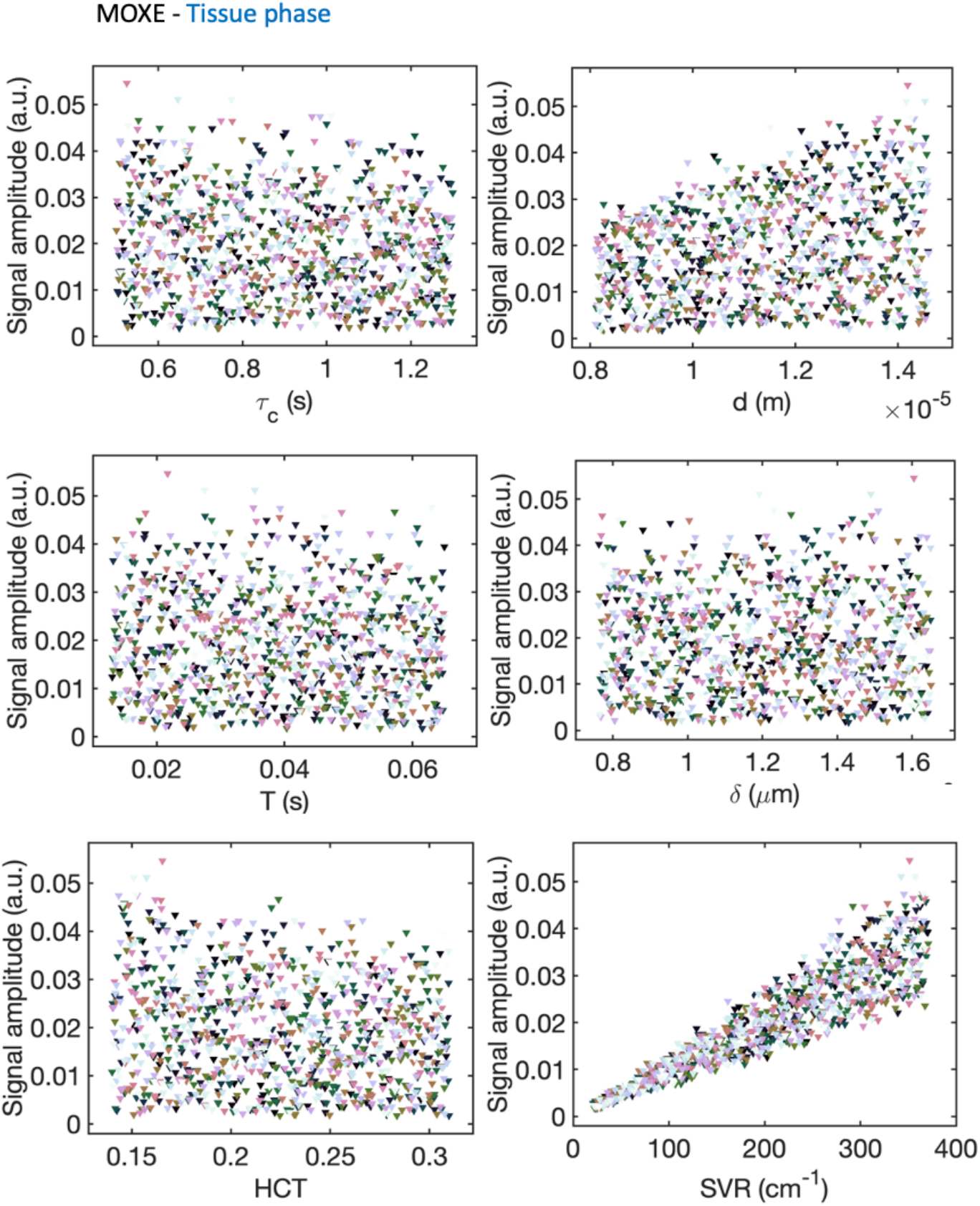
Scatter plots illustrate the signal relationship with each parameter individually while considering the complete variation of the other model parameters for the tissue phase in the MOXE model.

**Figure 9.**
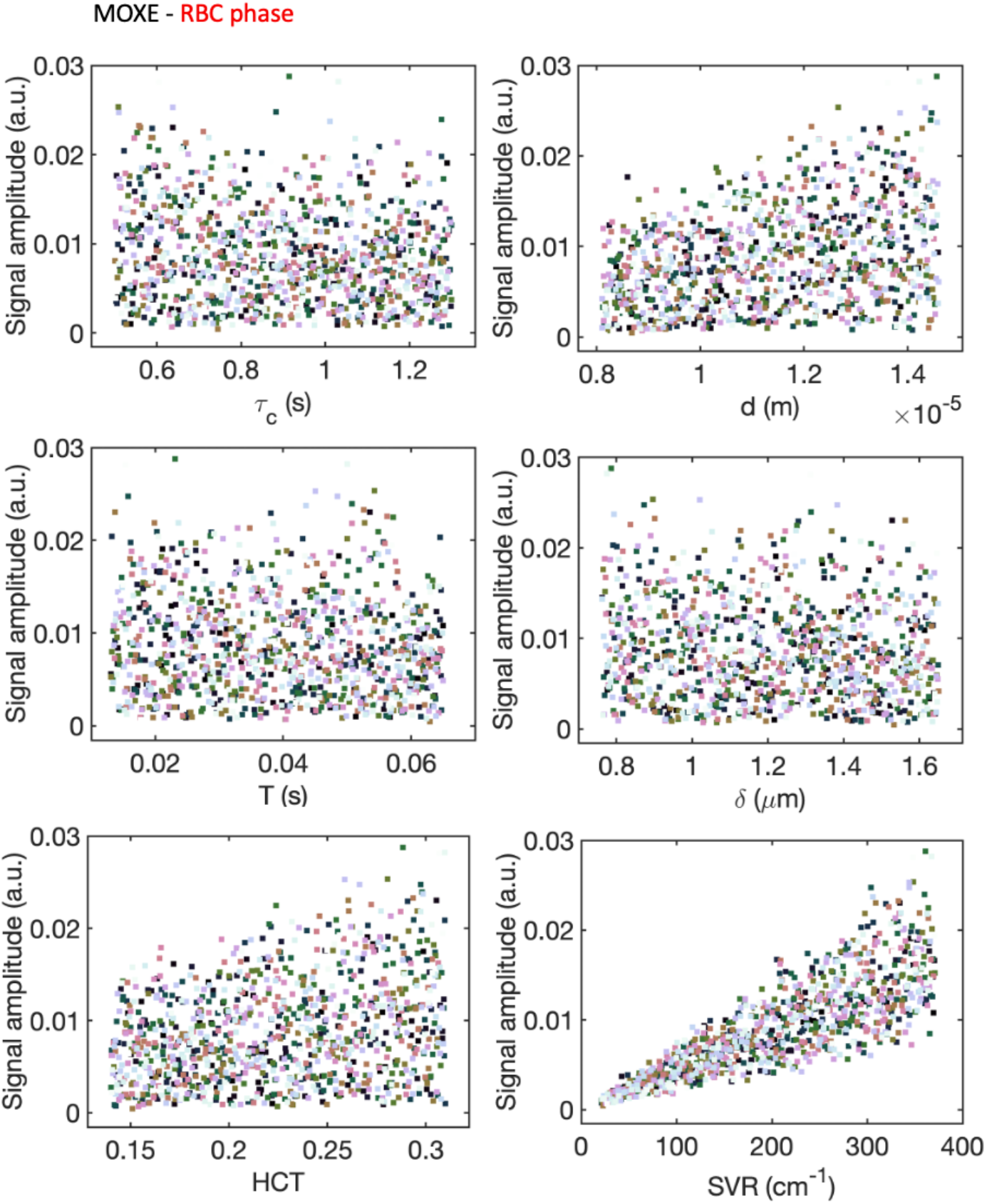
Scatter plots showing the signal relationship with each individual parameter whilst allowing the full variation of the other model parameters for the RBC phase in the MOXE model.

Sup. Figs. S1 and S2 show representative model sensitivity in a range of parameter combinations for the tissue and RBC phase respectively. Fifteen combinations were produced from the six-parameter MOXE model assessing two parameter interactions.

## Discussion

Univariate, scatter, and SPC sensitivity analysis methods were employed to investigate the Patz and MOXE models’ sensitivity to changes within their respective input parameter space.

### Univariate Sensitivity Analysis

Univariate sensitivity analysis simulations for the Patz model displayed the greatest sensitivity to SVR throughout the entire range of TR values. Sensitivity to *d* and τ_*c*_ was lower but increased with TR. The inverse relationship between τ_*c*_ and signal amplitude is apparent in Fig. 2 and is dependent on the proportion of ^129^Xe within the GEZ. The plug flow highlighted in Fig. 1 depicts the surface area apportioned to the individual regions traversing the GEZ. Following ^129^Xe diffusion into the septum (Patz) or capillary (MOXE), each region then reflects the fraction of blood in the GEZ occupied by ^129^Xe. At smaller τ_*c*_ values the proportion of ^129^Xe within the GEZ increases (Eq. 13), resulting in an increase in signal amplitude. This relationship is also reflected in both MOXE tissue and RBC phase τ_*c*_ simulations.

Univariate sensitivity analysis simulations for the MOXE model included assessments for both the tissue phase and RBC compartments. The tissue phase results reflect a combination of the ^129^Xe contribution in the blood plasma and lung tissue (Eq. 20), and therefore represent the larger overall pool of dissolved ^129^Xe. The RBC phase results reflect a comparatively smaller ^129^Xe concentration contribution (Eq. 21), highlighted by the reduced signal amplitudes in each parameter-specific plot (Fig. 4), relative to the plots for the tissue phase (Fig. 3).

The MOXE model parameters that are shared with the Patz model (SVR, d, τ_*c*_), displayed similar results when evaluating the model sensitivity within the tissue phase (Figs. 2 and 3). Both models exhibited an increase in model output range as a function of increasing TR due to the flow component associated with ^129^Xe reaching the blood after approximately 100 ms.

Increases in SVR, 5, *d* led to increases in tissue phase signal across the TR range, whereas HCT, *T* and τ_*c*_ showed inverse relationships demonstrating a reduction in the overall blood contribution. (Fig. 3). The opposite relationship was observed for HCT in the RBC phase, as HCT largely dictates the contribution of the ^129^Xe RBC signal within the GEZ (Eq. 20).

Plots for *δ* and *T* in the tissue phase (Fig. 3) show larger output ranges at shorter TRs, demonstrating a greater proportion of ^129^Xe within the tissue prior to entry into the GEZ. Conversely, longer TRs depict smaller ranges and reflect the dispersion of ^129^Xe away from the tissue and into the blood stream.

### Signal Percentage Change

Results for the SPC were analysed for each parameter from each model to determine if specific ranges within the acquisition window exhibit greater changes in sensitivities than others. Fig. 5 displays the SPC over the range of TR periods for the tissue phase and RBC within the MOXE model and for the dissolved phase within the Patz model. For the MOXE model, most of the variation in signal change due to HCT, *d*, and *T* occur at TR values < 200 ms within the tissue phase, while change due to *δ* and τ_*c*_ varies more slowly over the TR range. SVR displays the largest, and unchanging effect over the entire TR range. These results are in agreement with experimental validation of saturation recovery modelling techniques (19). Similar results were established within the RBC phase, with the exception of 5 which remained approximately constant apart from within the initial TR range (TR ≤ 100 ms).

SPC plots for the dissolved phase in the Patz model displayed greater variability for *d* than seen with the MOXE model, with variation again mainly limited to the TR < 200 ms range. τ_*c*_ and SVR vary in a similar way to the variation seen within the MOXE model.

### Scatter plots

Scatter plots offer an intuitive way to visualise the relationship between specific parameters and model outputs, which, unlike the univariate analysis, accounts for the influence of all other parameters. Scatter plots for the three Patz and six MOXE parameters yielded similar conclusions to corresponding model univariate SA methods. 2D plots for shared Patz and MOXE parameters (Figs. 6, 8, and 9) displayed high positive signal correlations with SVR that are relatively uncontaminated by the effects of other parameters. Plots of *d* and τ_*c*_ showed a degree of positive and negative correlation, respectively, but with far less specificity, as illustrated by the broad scatter of sample points. Plots for the MOXE-specific parameters *δ, T* showed little evidence of correlation, indicating low specificity of signal changes in both the tissue and RBC phases, whereas HCT demonstrated a greater degree of correlation, particularly in the RBC phase, indicating its stronger influence on the observed signals (Figs. 8 and 9).

### 3D plots

3D plots displayed interaction effects between the three Patz and six MOXE parameters, by assessing the increase or decrease in signal amplitude correlation as highlighted in the 3D surface plots (aerial perspective) in Fig. 7, Sup. Figs. S1 and S2. Interaction effects for the Patz model were evident between all parameters with stronger correlations in SVR - *d* and τ_*c*_ - *d* combinations and are heightened dependent on TR value, demonstrated by the τ_*c*700_ plots in Fig. 7. For MOXE tissue phase plots (Sup. Figs. S1 and S2), interactions were observed in all parameter couplings with less noticeable correlations observed when assessing *T*, and *δ*. Both parameters showed little model output variability within the univariate method and displayed weak correlations in the scatter. As such, minimal interactions with these parameters at the specific delay time chosen (∼400 ms) can be seen. Interactions between SVR and *d* and SVR and HCT displayed more noticeable interactions when evaluating the upper limits of SVR and *d*, against the lower limits of HCT. As the HCT is RBC phase-specific, as previously mentioned, lower values would ensure maximum tissue phase signal contributions for SVR and *d*. Similarly, noticeable interactions were observed for all MOXE RBC phase parameter couplings. The largest interactions were observed in SVR against HCT, and SVR against *d*. The lowest interactions were observed with SVR against T, *δ*, and τ_*c*_. However, unlike tissue phase simulations, RBC simulations displayed an increased sensitivity when assessing the range of 5, resulting in stronger correlations in comparison to the tissue phase. 3D plots for the RBC compartment for SVR, *d* and τ_*c*_ interactions displayed similar trends to the 3D scatter Patz plots (Sup. Figs. S1 and S2).

### Implications for *in vivo* measurements and limitations

Sensitivity varied for the different models and dissolved phase compartments within MOXE, but the surface area to volume ratio (SVR) and septal wall thickness (*d*) consistently demonstrated greater sensitivity in both the Patz and MOXE models. The alveolar-capillary barrier thickness (*δ*) showed the greatest difference in sensitivity between the two MOXE model dissolved phases, but overall, sensitivity to variation in *δ* is low, which has previously been demonstrated experimentally (19). Sensitivity to all other parameters was marginally greater than *δ*, but substantially lower than sensitivity to SVR and *d*. These observations provide useful guidance for the likely signal-to-noise ratio (SNR) requirements of CSSR measurements designed to measure model parameters. For example, an SNR of only ∼2 would be adequate to distinguish the extremes of the range of possible SVR and *d*, whereas an SNR of ∼15 would be required to distinguish the extremes of the range of possible *δ* (Figs. 3 and 4). More subtle differences would require correspondingly higher SNR.

Parameter ranges chosen for each model were extracted from a limited set of sources and measurement techniques (Table 1). As a result, the ranges highlighted for each parameter may possess intrinsic biases. Moreover, a broad range of both healthy and diseased values was used for each sensitivity analysis technique without differentiating between disease type and stage of progression. The analysis of stratified disease groups for both models would permit the evaluation of model sensitivity to disease classification and progression.

Both models demonstrate higher sensitivity to TR measurements between 0 - 200 ms. Different studies in the literature have evaluated ^129^Xe signals over different TR ranges. For example, (6; 22) assessed the Patz and MOXE models at TR ≤ 200 ms, with a range of TR measurements (TR 200 < ms and TR > 200 ms) being more common generally in other studies (4; 19; 15). These results suggest that future studies would benefit from performing comparable simulations before finalising their experimental protocols.

It is possible that further sensitivity analyses may yield additional useful information. The main objectives of the presented sensitivity analysis of the Patz and MOXE models were parameter ranking and screening; however, mapping - which observes areas within the parameter input space producing extreme output values - was not evaluated. The inclusion of mapping methods such as the distance-based generalised sensitivity analysis (DGSA) (10) in conjunction with the prior methods may provide a further understanding of model sensitivity.

## Conclusions

The sensitivity analysis of the Patz model demonstrated a higher signal sensitivity to both SVR and *d* than to τ_*c*_. Similarly, the sensitivity analysis of the MOXE model also displayed sensitivity to SVR and *d*, whilst also demonstrating sensitivity to HCT and *T*. In contrast, a much lower sensitivity to 5 was identified. Sensitivity variation as a result of delay time increased between the first 200 ms for both models due mainly to the underlying processes surrounding saturation of the tissue and blood flow. Consequently, multiple measurements taken within periods between 0 - 200 ms were highlighted as advantageous in optimising model sensitivity due to the intrinsic parameter variation highlighted in both the signal percent change and scatter plot analyses. Interaction effects within the Patz and MOXE model were also shown to have an effect on the likely specificity of changes in signal to changes in the underlying parameters. These findings demonstrate the likely ability of two hyperpolarised ^129^Xe diffusion models to identify differences in lung microstructure and function.

## Acknowledgments

This work is co-funded by an EPSRC Industrial CASE award (Voucher No. V20000074) aligned to the EPSRC UCL Centre for Doctoral Training in Medical Imaging (EP/S021930/1) and Glaxo-SmithKline Research Development Ltd (BIDS3000035683).

## Figures and Tables

**Figure S1:**
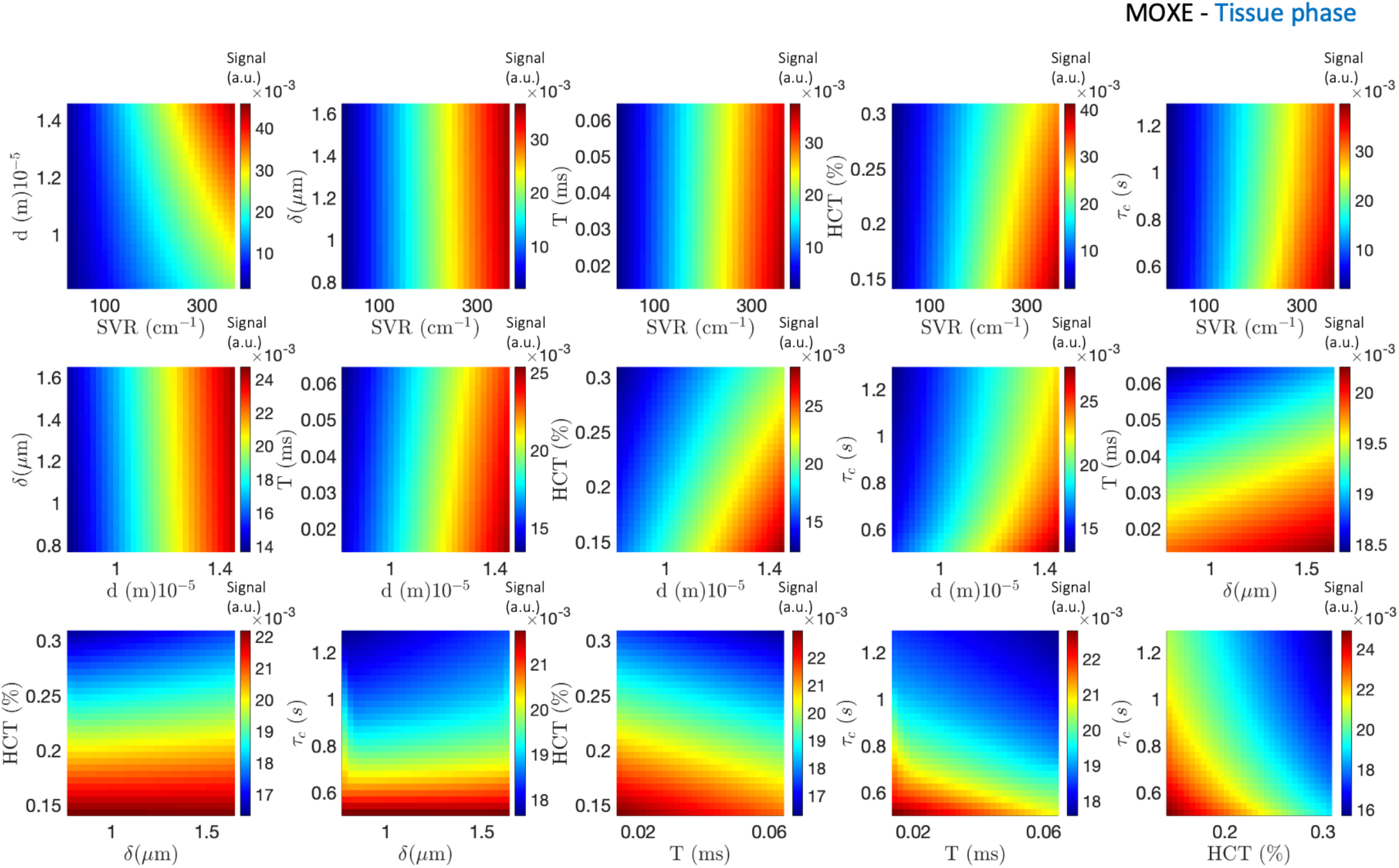
3D plots show the relationship between interaction effects of all other parameters in the TP of the MOXE model. All simulations were taken at a TR of 400 ms. The colour bar indicates signal intensity, where the direction of the trend corresponds to the contribution of interaction.

**Figure S2:**
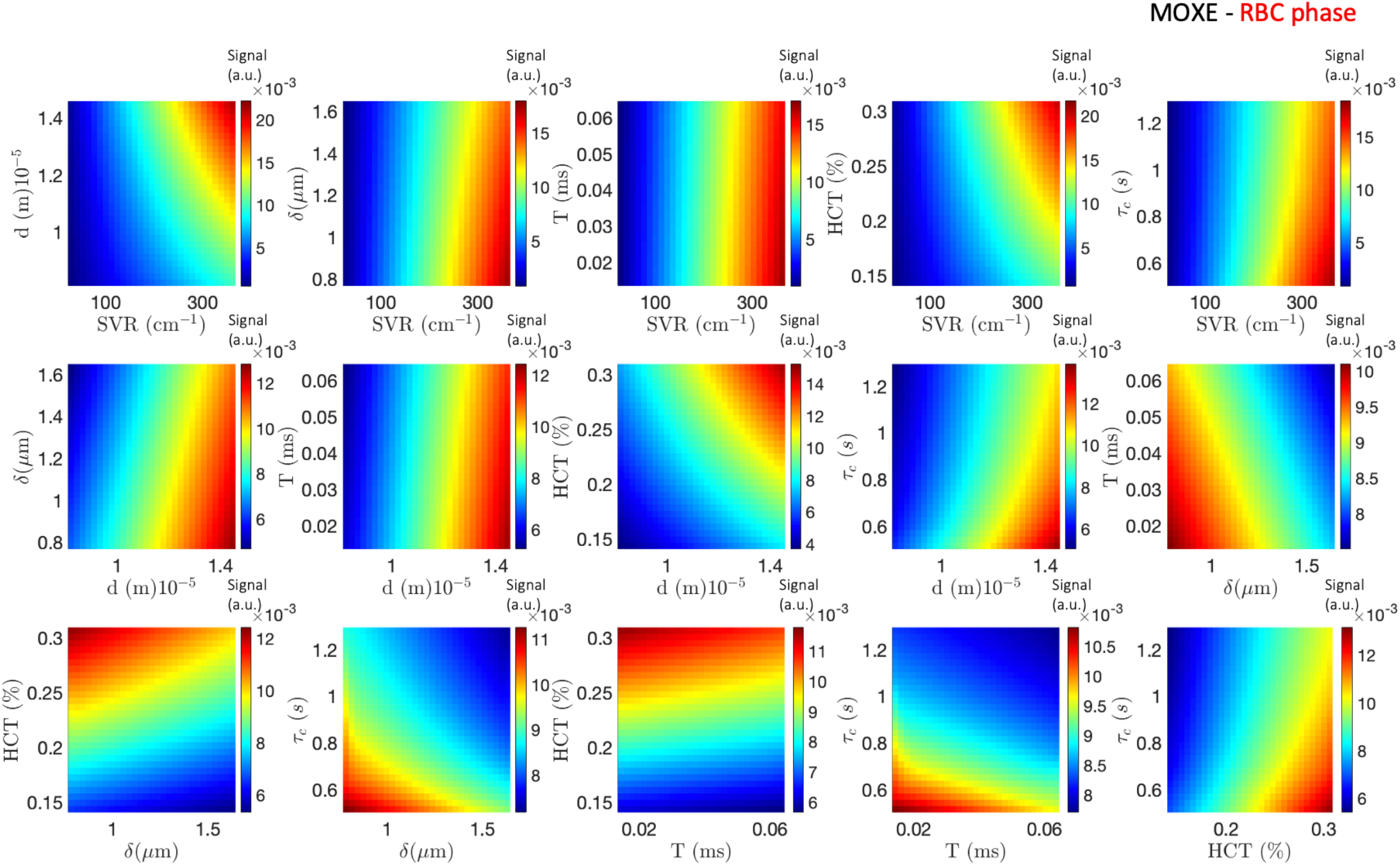
3D plots show the relationship between interaction effects of all other parameters in the RBC of the MOXE model. All simulations were taken at a TR of 400 ms. The colour bar indicates signal intensity, where the direction of the trend corresponds to the contribution of interaction.

